# Atypical Visual Selective Attention in Children with Dyslexia: Evidence from N2pc and PD

**DOI:** 10.1101/2024.12.24.630007

**Authors:** Hongyu Liu, Yulu Sun, Zimo Wang, Jialiang Guo, Yan Song, Xiangzhi Meng

## Abstract

Efficient visual attention is fundamental to the development of reading abilities. Previous studies have identified visual attention impairments in individuals with dyslexia, particularly in visual attention span and sluggish attention shifting. However, findings regarding basic visual selective attention remain controversial. To address this issue, the present study provides event-related potential (ERP) evidence to verify whether children with developmental dyslexia (DD) suffered from deficits in visual selective attention. A pop-out visual search task was used to examine visual attentional patterns to a lateral target in 66 children (33 children with DD and 33 typically developing (TD) children) using electroencephalography. Compared to TD children, children with DD showed a larger and prolonged P1 component, as well as a larger target-evoked N2pc component. The larger P1 amplitude was associated with lower reading fluency while the larger N2pc amplitude was associated with poor reading comprehension performance. Interestingly, the target-elicited N2pc was followed by positivity (P_D_ component) in TD children but not in children with DD; however, the N2pc amplitude was correlated with the P_D_ amplitude in children with DD. Children with DD show immature early attentional processing and may have an imbalance in the regulation of visual selective attention between attentional selection and suppression. Our findings provide ERP evidence for interpreting the underlying neural mechanism of visual selective attention in children with DD and may offer a new perspective for exploring the mechanism of visual attention deficits in dyslexia.

**Research Highlights:**

- In a pop-out visual search task, children with dyslexia exhibited a larger N2pc and an absent P_D_ component.
- The absence of the P_D_ component in children with dyslexia may be associated with their larger N2pc amplitudes.
- N2pc amplitude is associated with reading ability in children with dyslexia.
- Children with dyslexia need to allocate more resources towards the target but also struggle to disengage from the attended location.

## Introduction

Successful reading is inseparable from efficient visual attention, which enhances objects recognition, reduces interference from irrelevant information, and facilitates word encoding and memory storage (Vecera & Rizzo, 2003). In particular, reading, as a dynamic process, requires normal visual attention to ensure successive eye movements and quick switching across words and sentences (Vidyasagar & Pammer, 2010). Thus, visual attention plays a fundamental role in effective reading and learning.

Given the importance of visual attention in reading, deficits in visual attention may lead to developmental dyslexia, a neurodevelopmental disorder characterised by difficulties in accurate reading, fluent word recognition and spelling (American Psychiatric Association, 2013). For years, the core deficits in dyslexia have been debated, particularly phonological and visual impairments. Numerous studies have argued that the main causes of dyslexia are phonological and linguistic deficits(for a review, see Peterson & Pennington, 2015), leading to difficulties in the representation, storage and extraction of phonetics, which in turn results in poor reading abilities (Vellutino, Fletcher, Snowling, & Scanlon, 2004). However, phonological deficit theory fails to fully explain dyslexia, especially in non-linguistic tasks (Castles & Coltheart, 2004; Hokken, Krabbendam, van der Zee, & Kooiker, 2023; Nguyen et al., 2021). In recent years, a growing body of evidence suggests that it is visual attention impairments that contribute to dyslexia, rather than phonological deficits (Gabrieli & Norton, 2012; Vidyasagar et al., 2010; White, Boynton, & Yeatman, 2019). Such visual attention impairments, primarily deficits in the visual span and sluggish attentional shifting, have been observed in both children with dyslexia and those with familial risk (Facoetti, Corradi, Ruffino, Gori, & Zorzi, 2010; Lallier et al., 2010) and do harm to reading abilities (Ebrahimi, Pouretemad, Stein, Alizadeh, & Khatibi, 2022; Franceschini, Gori, Ruffino, Pedrolli, & Facoetti, 2012; Gavril, Roșan, & Szamosközi, 2021; White et al., 2019). However, these theories reveal the difficulties in integrating multiple attentional processes into visual attention, making it difficult to isolate specific visual attentional processes in dyslexia. To address issue, it is essential to investigate more fundamental attentional mechanisms, such as visual selective attention.

Visual selective attention encompasses maintaining attention on relevant information among numerous irrelevant distractors (Carrasco, 2011) and can be measured using feature search tasks. Understanding the mechanism of visual selective attention during a visual search may reveal how foundational visual attentional processing works in dyslexia, as it excludes the influence of working memory or higher-order executive processes which are impaired in individuals with dyslexia (Daucourt, Schatschneider, Connor, Al Otaiba, & Hart, 2018; Garcia, Mammarella, Tripodi, & Cornoldi, 2014; Gathercole, & Pickering, 2000). Previous studies have found that the behavioural performance of readers with dyslexia was comparable to that of their typically developing peers in simple feature search tasks, but slower in conjunction search tasks when more distractors were present (Iles, Walsh, & Richardson, 2000; Nguyen et al., 2021; Roach, & Hogben, 2007). These results suggest that readers with dyslexia have no problems with efficient feature search tasks, but rather show inefficiencies in tasks with heavy attentional loads. However, some studies have reported slower visual searches even in simple feature tasks among individuals with dyslexia (Jones, Branigan, & Kelly, 2008; Sigurdardottir, Omarsdottir, & Valgeirsdottir, 2024), arguing that readers with dyslexia may struggle with precise visual stimulus localisation. Therefore, it remains unclear whether individuals with dyslexia have difficulties with visual selective attention. These mixed results may stem from the interplay between bottom-up attentional capture (Theeuwes, 1991) and top-down control (Ruff & Driver, 2006), as simple feature searches rely primarily on the former, whereas conjunction searches depend more on the latter. Furthermore, previous studies have uncovered differences in attentional functioning that are not captured by behavioural measures. In summary, direct neurophysiological evidence regarding whether dyslexia is associated with deficits in selective attention is lacking. Accordingly, the present study investigated the neural mechanisms of visual selective attention in children with dyslexia as a potential contributor to reading difficulties.

Electroencephalography (EEG) provides high temporal resolution, enabling the examination of neural responses over time during a visual search. The N2-posterior contralateral (N2pc), a negative component of the visual cortex contralateral to the attended target, reflects visual selective attention (Luck & Hillyard, 1994; Stoletniy, Alekseeva, Babenko, Anokhina, & Yavna, 2022). N2pc is a reliable indicator for both adults and typically developing children (Sun et al., 2018) and has been used to classify children with attention deficit/hyperactivity disorder (ADHD) who suffer from attentional deficits (Li et al., 2023; Wang et al., 2016). Thus, the N2pc is a powerful index for assessing selective attention ability and identifying individuals with general attentional deficits (Hu, Tang, & Huang, 2023; Li et al., 2023). To the best of our knowledge, only one recent study has investigated the N2pc component in children with dyslexia using an object substitution masking task (Koffman, Flaten, Desroches, & Kruk, 2023). Although children with dyslexia showed a general trend towards greater N2pc than the control groups, this effect was not significant. P_D_, the positive component contralateral to a suppressed item, reflects the active suppression of distractors (Hickey, Di Lollo, & McDonald, 2009) and the termination of attention to an attended object (Jannati, Gaspar, & McDonald, 2013; Sawaki, Geng, & Luck, 2012). Previous studies have shown that dyslexics have difficulty in disengaging attention from the currently attended location (Lallier et al., 2010), which might be implicated in the P_D_ component. Therefore, the N2pc and P_D_ components may serve as markers for deficiencies and the underlying neural mechanisms of visual selective attention in children with dyslexia.

This study used a classic pop-out visual search task (Wang et al., 2016): 1) to investigate whether children with dyslexia exhibit impairments in covert visual selective attention as indexed by N2pc and P_D_, and 2) if so, to characterise whether visual selective attention ability is correlated with reading ability in children with dyslexia. Our study not only reveals the neural characteristics of visual selective attention in dyslexia, but also provides neural evidence supporting that the view that dyslexia arises from deficits beyond linguistic factors.

## Methods

### Participants

Seventy-two Chinese children aged 8-13 years participated in the experiment, including 36 children with developmental dyslexia (DD group) and 36 typically developing children (TD group). All participants were right-handed, had normal or corrected-to-normal vision, and an IQ of at least 90. One participant who was diagnosed with ADHD based on parental questionnaires was excluded from the study. Four participants with low task accuracy and one with poor data quality due to excessive muscle activity were also excluded. None of the remaining participants had a history of ADHD or other mental illnesses, according to parental questionnaires. Ultimately, our data for analysis was collected from 33 children in the DD group (17 females, mean age: 11.0) and 33 children in the TD group (13 females, mean age: 10.9) (**Table 1**).

**Table 1.** Demographic information of the children.

|  | DD group<br>(N=33) | TD group<br>(N=33) | Statistics |
| --- | --- | --- | --- |
| Age (years) | 11.0 ± 1.1 | 10.9 ± 1.0 | $t(64) = -0.388, p = .699$ |
| Gender (male : female) | 16 :17 | 20:13 | $\chi^2(1) = .978, p = .323$ |
| IQ | 114.0 ± 13.6 | 119.7 ± 9.9 | $t(58.4) = 1.981, p = .052$ |
| RAN times (s) | 15.0 ± 3.2 | 13.0 ± 2.8 | $T(64) = -2.771, p = .007^{**}$ |
| Chinese Character Reading <sup>a</sup> | 71.4 ± 21.6 | 104.1 ± 7.3 <sup>c</sup> | $t(45) = 5.510, p < .001^{***}$ |
| Character Reading Fluency <sup>a</sup> | 82.0 ± 11.5 <sup>b</sup> | / | / |
| CCPM <sup>a</sup> | 84.6 ± 8.3 <sup>b</sup> | / | / |
| Reading Comprehension <sup>a</sup> | 104.3 ± 8.8 <sup>b</sup> | / | / |

Note: <sup>a</sup> Values are standard scores to a 100-point scale according to the corresponding norms and standard deviation.
<sup>b</sup> Twenty-two DD children took part in these tests.
<sup>c</sup> Fourteen TD children took part in these tests.
RAN = rapid automatized naming; CCPM = character read correctly per minute; DD = developmental dyslexia; TD = typically developed.

Children with dyslexia were previously diagnosed by qualified psychologists and characterised using parental questionnaires. They were reassessed using the Chinese Character Reading Test and Character Reading Fluency Test. Both tests indicated standard scores below 90, with an average level of 100. Among the children with dyslexia, twenty-two took part in Character Reading Fluency and Reading Comprehension Test, as shown in **Table 1**. All children in the TD group achieved excellent scores on school reading tests and reported no difficulties in learning Chinese. Fourteen participated in the Chinese Character Reading Test and scored above the average level of 100, as shown in Table 1. No significant differences were found in age, sex, or IQ between the groups. The DD group showed significantly slower Rapid Automatized Naming (RAN) times (*t*(64) = −2.771, *p* = .007, Cohen’s *d* = .682) and poorer Chinese character reading performance (*t*(45) = 5.510, *p* < .001, Cohen’s *d* = 1.757) than the TD group. This study was approved by the Ethics Committees of Peking University and Beijing Normal University. Written consent was obtained from each child and their parents in accordance with the Declaration of Helsinki.

### Behavioural Tests

Five behavioural tests were used to assess reading-related skills, and the reading scores were standardised to a 100-point scale according to the corresponding norms and standard deviations.

#### Rapid Automatized Naming Test

The RAN test was employed to measure the children’s abilities in information processing and speech representation extraction (Georgiou, Parrila, Cui, & Papadopoulos, 2013). The children were required to name a matrix of digits (1-9 randomly arranged) as quickly and accurately as possible. The task was performed twice and the average reading time was used as the final score. Cronbach’s alpha for the task was 0.88.

#### Chinese Character Reading Test

The Chinese Character Reading Test, involving 150 characters, was used to test the children’s lexical reading accuracy (Meng & Lai, 2024). The children were instructed to read the characters aloud as accurately as possible without time limitations. The test was terminated when participants made 15 consecutive errors. The final score was recorded as the total number of characters correctly read. Cronbach’s alpha for the task was 0.96.

#### Character Reading Fluency Test

The children were presented with 160 Chinese characters, selected from first grade textbooks and arranged in a matrix in ascending order of difficulty (Meng et al., 2024). They were instructed to read out the characters as quickly as possible within one minute. Higher scores reflected greater fluency in character recognition. Cronbach’s alpha for the task was 0.97.

#### Reading Comprehension Test

To evaluate the children’s text reading fluency and comprehension, we administered the Chinese Passage Reading Test (Meng et al., 2024). Passages of varying difficulty levels were used depending on the children’s grades. They were asked to read out the passage, both accurately and quickly. Upon completion, they were tested using eight questions on the content of the passages to evaluate their comprehension. We quantified reading fluency using the Character Read Correctly Per Minute (CCPM), which was derived by dividing the total number of characters read correctly by the individual’s reading time, in seconds. Simultaneously, questions answered correctly served as an indicator of reading comprehension. Cronbach’s alpha was 0.94 and 0.73 for the CCPM and reading comprehension task, respectively.

### Stimuli and Design in EEG task

The visual search task is the typical paradigm used in previous studies (Li et al., 2023; Sun et al., 2018; Wang et al., 2016). In this study, the stimulus was a circular search array comprising one filled circle as the target and 11 filled diamonds as distractors. Each item was positioned at a visual angle of 5° from the central fixation point (**Figure 1A**). The target circle appeared randomly in either the right (2 or 4 o’clock position) or left (8 or 10 o’clock position) hemifield during each trial. The fixation cross was presented for a jittered interval ranging from 900 to 1100 ms, followed by a circular search array for 200 ms. A blank screen was then displayed for up to 2800 ms, which disappeared when the participants responded by pressing a key.

**Figure 1.**
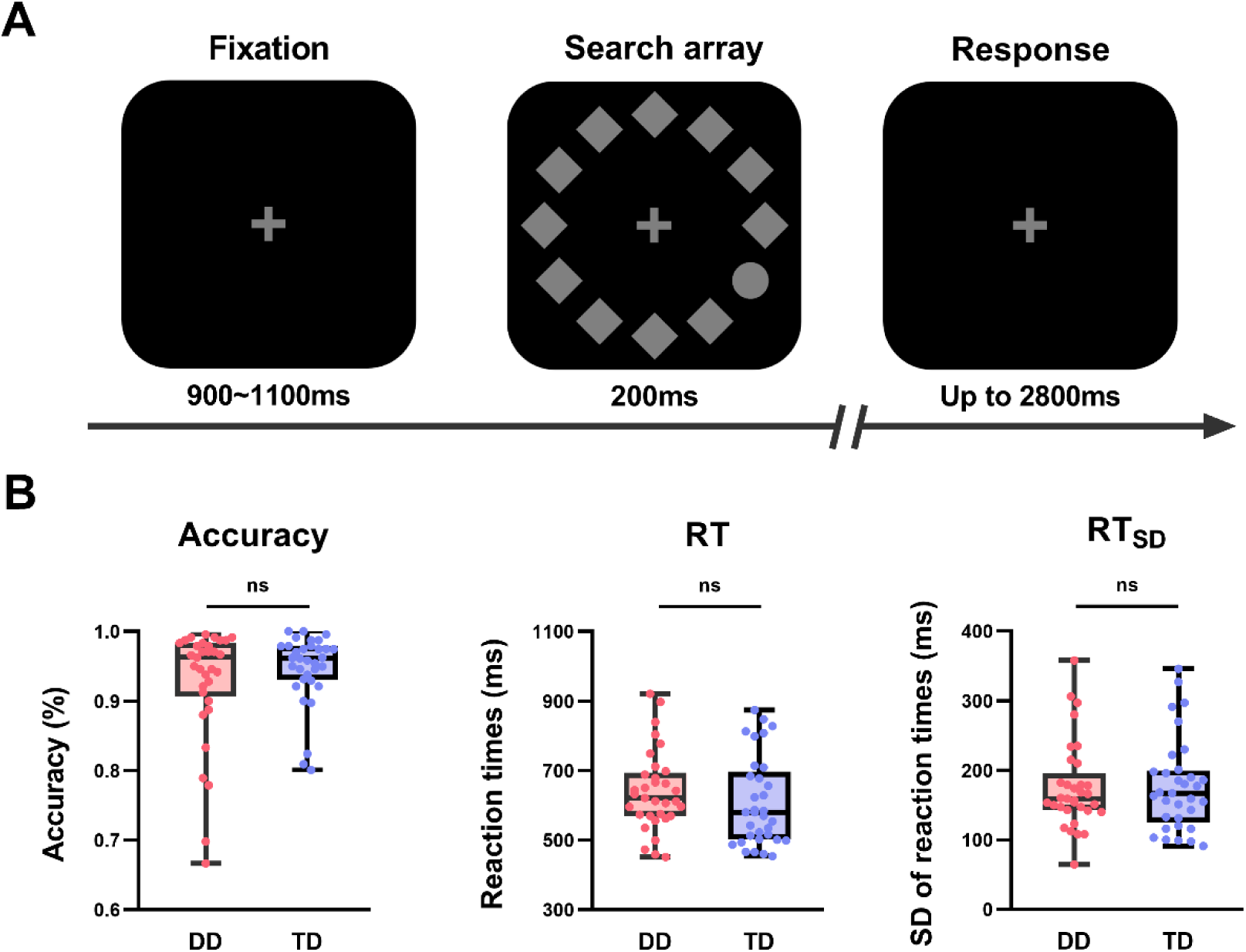
Trial sequence and behavioural results. **(A)** An example of the trial displays. The search array stimulus consisted of one circle as the target and 11 diamonds as distractors. **(B)** The behavioural results of the experiment including accuracy (ACC), reaction times (RT) and standard deviation of reaction times (RTSD) in DD and TD groups.

The participants were instructed to maintain their gaze on the fixation cross throughout the experiment and to report whether the target appeared in the upper or lower visual field as quickly and accurately as possible. The formal experiment consisted of eight blocks, each containing 30 trials. Prior to the formal experiment, a practice session was conducted to ensure that the participants understood the task properly.

### Behavioural analysis

Only trials in which the participants responded correctly were included in the analysis. Reaction time (RT) was calculated from the onset of the stimuli. Reaction time standard deviation (RTSD) was used to quantify the variability in response times across trials.

### EEG recording and analysis

During the visual search task, EEG data were recorded using a 128-channel HydroCel Geodesic Sensor Net (Electrical Geodesics, Inc., Eugene, OR, USA) with Net Station EEG Software. Electrode impedance was kept below 50 kΩ during acquisition. Data were digitised at 1000 Hz and bandpass filtered at 0.01–400 Hz with Cz as the online reference.

The raw EEG data were processed using standard functions from the EEGLAB toolbox (Delorme & Makeig, 2004) in MATLAB. Data from 33 lateral electrodes were removed due to susceptibility to eye, face and head movements. EEG data were then resampled to 250 Hz, bandpass filtered at 1–30 Hz using a zero-phase finite impulse response (FIR) filter and re-referenced to the average of all remaining electrodes. Electrodes containing more than 5 seconds of artefacts or high-frequency noise deviates by 4 standard deviations were interpolated using a spherical spline interpolation and excess artefacts were manually removed. Independent component analysis (ICA) was used to remove components related to eye movements, heartbeats, and head movements. For each trial, the EEG data were segmented to stimulus onset (−200 to 600 ms), and the baseline (−200 to 0 ms) was subtracted. Epochs with blinks at Fp1 and Fp2 electrodes exceeding ±75 μV, horizontal eye movements at F9 and F10 electrodes exceeding ±50 μV, and voltages exceeding ±100 μV were automatically rejected. Trials with incorrect responses and response times over 2000 ms or shorter than 200 ms were also excluded. Of the 167 and 191 remaining epochs in the DD and TD groups (*t*(64) = 2.788, *p* = .007, Cohen’s *d* = 0.686), respectively, 80 epochs were randomly selected to eliminate the effects of different trial numbers between the two groups.

Based on the visual inspection of topographic maps and previous studies, event-related potentials (ERPs) at PO7 and PO8 were calculated for the contralateral and ipsilateral waveforms to the target. A 20 ms window around the maximum value in the group-averaged ERP was used to obtain the individual peak values and peak latencies for P1 at 80-150 ms. The N2pc (200-300 ms) and P_D_ (300-400 ms) components were derived from the difference waves, subtracting ERP waveforms at ipsilateral electrodes from the waveforms at contralateral electrodes to the target. Then each participant’s amplitudes were measured using a 20 ms window around the peak value in the difference waves. The onset latencies of the N2pc and P_D_ were estimated using the Jack-knifing method for the time at which each component reached 50% of its peak amplitude (Miller, Patterson, & Ulrich, 1998).

For statistical analyses, one-sample t-tests were used to examine whether the amplitude of each component differed significantly from zero. The difference in the P1 component between groups was measured using a two-way mixed-measures ANOVA, with Group (DD, TD) as the between-subject factor and Lateralisation (contralateral, ipsilateral) as the within-subject factor. If the Group × Lateralisation interaction effect reached significance, the Holm–Bonferroni correction was applied for simple effect analysis. For the N2pc and P_D_ components, to accurately measure their amplitudes while avoiding bias from fixed time windows, we applied a nonparametric permutation approach (Sawaki et al., 2012) to confirm that the statistical results of the N2pc and P_D_ components did not result from chance. We randomly labelled the side of the target appearance for each trial, re-averaged the data and calculated the area values from the difference waveforms between 200 and 400 ms. This procedure was repeated 1000 times to obtain a null distribution of the area values. If the actual area exceeded 95% of the randomised values, we reject the null hypothesis that the observed N2pc or P_D_ was merely a product of random noise.

To examine the relationship between N2pc and P_D_ components, as well as the associations between ERP responses and reading abilities, we performed a partial Pearson correlation analysis controlling for age and IQ to exclude the impact of development and other underlying factors.

## Results

### Behavioural results

As illustrated in **Figure 1B**, no significant difference in accuracy was found between the children in the DD and TD groups (*t*(64) = 1.150, *p* = .254, Cohen’s *d* = 0.283). Also, the two groups did not show a significant difference in RT (*t*(64) = −1.072, *p* = .288, Cohen’s *d* = 0.264) or RTSD (*t*(64) = 0.123, *p* = .903, Cohen’s *d* = 0.03). Thus, the DD group completed the visual search task as accurately and quickly as the TD group.

### ERP results

#### P1

As **Figure 2A,B** showed, a reliable P1 component was elicited in the contralateral and ipsilateral hemispheres after stimulus onset in the DD group (Contralateral: *t*(32) = 9.987, *p* < .001, Cohen’s *d* = 1.739; Ipsilateral: *t*(32) = 9.657, *p* < .001, Cohen’s *d* = 1.681) and the TD group (Contralateral: *t*(32) = 8.895, *p* < .001, Cohen’s *d* = 1.548; Ipsilateral: *t*(32) = 9.618, *p* < .001, Cohen’s *d* = 1.674). A two-way mixed ANOVA revealed that the P1 amplitude was significantly larger in the DD group than in the TD group (Group effect: *F*(1,64) = 5.186, *p* = .026; Group × Lateralisation interaction effect: *F*(1,64) = 0.077, *p* = .782). Moreover, P1 peak latency was significantly prolonged by approximately 10 ms in the DD group compared to the TD group (approximately 118 ms for the DD group; approximately 108 ms for the TD group; Group effect: *F*(1,64) = 11.951, *p* < .001; Group × Lateralisation interaction effect: *F*(1,64) = 0.971, *p* = .328; **Figure 3**). No significant Lateralisation effect was found for P1 amplitude (*F*(1,64) = 0.096, *p* = .757) or P1 latency (*F*(1,64) = 2.698, *p* = .105).

**Figure 2.**
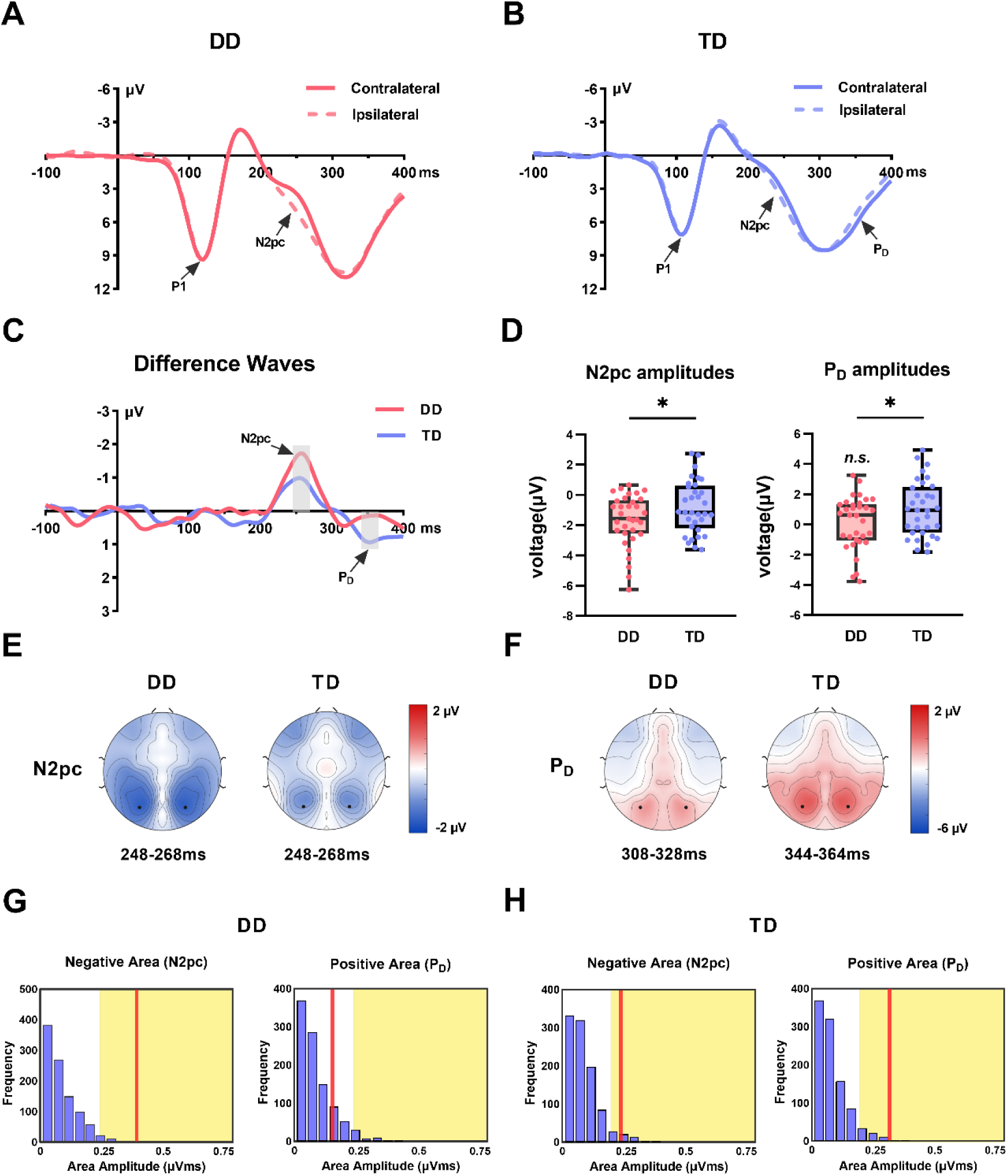
ERP results and topographic map distribution. Averaged ERPs at contralateral and ipsilateral electrodes to the target from PO7/8 in DD group **(A)** and TD group **(B)**. **(C)** Grand averaged difference waveforms obtained by subtracting ipsilateral from contralateral waveforms. The areas highlighted in grey show the significant window for the analysed components. The averaged N2pc and P_D_ amplitudes **(D)** and the topographic distribution for the N2pc **(E)** and P_D_ **(F)** components in each group. The black dots were the electrodes (PO7/8) used for analysis. Results of permutation tests on the N2pc negative and P_D_ positive areas from 200 to 400 ms were presented separately for the DD group **(G)** and TD group **(H)**. The purple bars indicated the estimated null distribution from 1000 permutations. The yellow areas represent the top 5% of the permutation distribution. If the red lines fall into the yellow areas, the observed values are significantly above the level of random noise. * *p* < .05; DD = developmental dyslexia; TD = typically developing.

**Figure 3.**
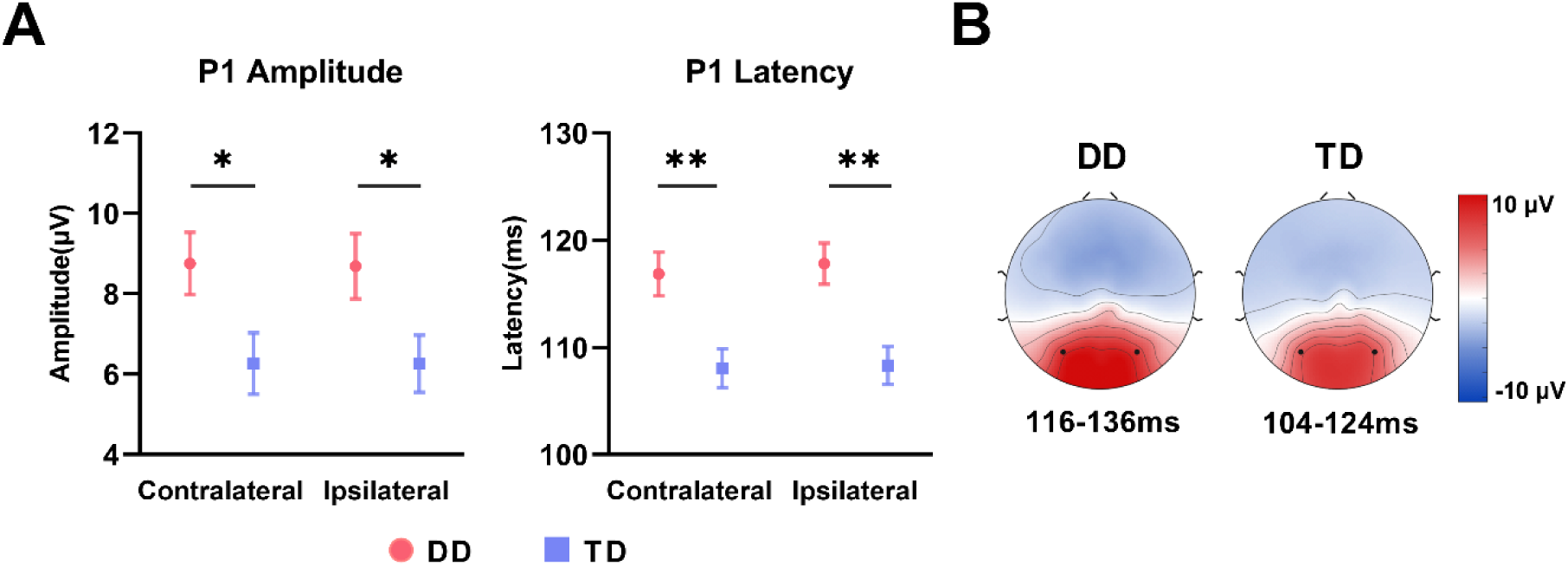
Group comparison of the P1 component. **(A)** The statistical results for the amplitude and peak latency of P1 component at the electrodes contralateral and ipsilateral to the targets at PO7/8. **(B)** Topographical distribution of P1 component. The black dots were the electrodes (PO7/8) used for analysis. * *p* < .05; ** *p* < .01; DD = developmental dyslexia; TD = typically developing.

#### N2pc

The target elicited a contralateral minus ipsilateral lateralisation effect at PO7/8 after stimulus onset for 200-300 ms in both groups **(Figure 2C,E**), suggesting that the lateral target induced the N2pc component in both the DD and TD groups. One sample t-tests showed that the N2pc (246-266 ms) was significantly different from zero in both the DD (*t*(32) = −5.610, *p* < .001, Cohen’s *d* = 0.977) and TD groups (*t*(32) = −2.666, *p* = .012, Cohen’s *d* = 0.464). Permutation tests confirmed this result (**Figure 2G,H**). An independent-samples t-test revealed that the N2pc amplitude was significantly larger in the DD group than that in the TD group (*t*(64) = 2.071, *p* = .042, Cohen’s *d* = 0.510, **Figure 2D**). No significant difference was found in the N2pc onset latency between the two groups (*t*(64) = 2.775, *p* = .106, Cohen’s *d* = 0.404).

#### P_D_

The P_D_ component, characterised by a positive contralateral-minus-ipsilateral lateralisation at PO7/8, 300-400 ms after stimulus onset following the N2pc component, was present in the TD group but not in the DD group (**Figure 2C,F**). One sample t-tests confirmed that the P_D_ component (342-362 ms) was significantly different from zero in the TD group (*t*(32) = 2.616, *p* = .013, Cohen’s *d* = .455), indicating the clear presence of the P_D_ component. In contrast, the P_D_ amplitude (308-328 ms) did not differ significantly from zero in the DD group (*t(32)* = 0.955, *p* = .347, Cohen’s *d* = 0.166). Permutation tests confirmed this result (**Figure 2G,H**). An Independent sample t-test indicated that the P_D_ component in the TD group was significantly larger than that in the DD group (*t*(64) = 2.142, *p* = .036, Cohen’s *d* = 0.527; **Figure 2D**).

### Correlation

#### Relationship between N2pc and P_D_

Compared with the TD group, the DD group showed a larger N2pc component but lacked a P_D_ component. It appears that children with dyslexia deploy insufficient attentional suppression because they over-allocate attentional selection resources to processing the pop-out targets. To test this hypothesis, we performed partial Pearson correlation analyses between N2pc and P_D_ amplitudes in the two groups separately, controlling for age and IQ. Surprisingly, there was a significant positive correlation in the DD group (*r* = 0.579, *p* < .001), but not in the TD group (*r* = 0.218, *p* = .223; **Figure 4A,B**). That means the larger the N2pc amplitude, the smaller the P_D_ amplitude. To minimise the potential bias induced by different time window selections between the two groups, we replicated the analysis on the P_D_ amplitude in the TD group using the same time window (308-328 ms) as in the DD group. Again, no significant correlation was found between the N2pc and P_D_ amplitudes in the TD group (*r* = 0.212, *p* = .253). Therefore, a significant correlation between the N2pc and P_D_ amplitudes was observed only in the DD group and not in the TD group.

**Figure 4.**
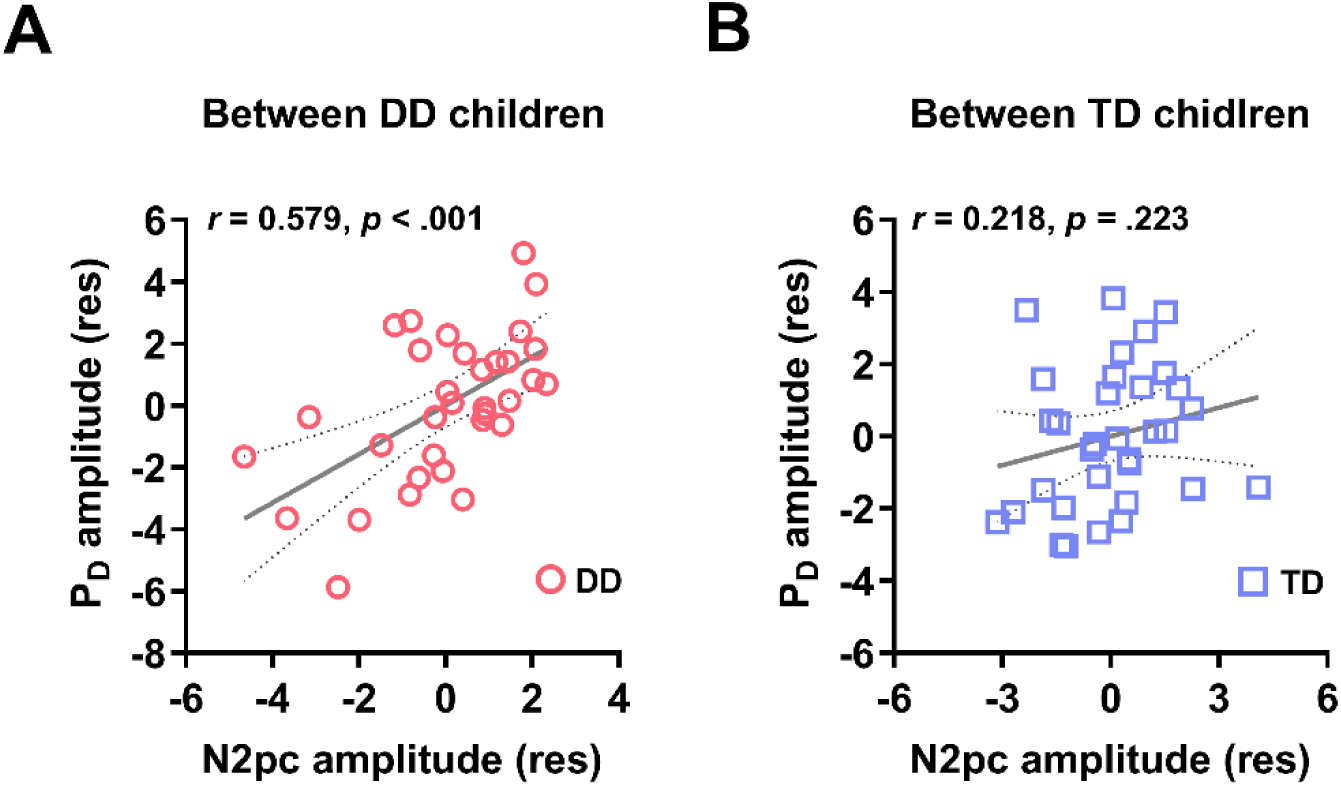
Relationship results between N2pc amplitudes and P_D_ amplitudes. Partial correlation results between the N2pc amplitudes and P_D_ amplitudes between DD children **(A)** and TD children **(B)**. The residuals of N2pc and P_D_ amplitudes after excluding the effects of age and IQ were used here. *** *p* < .001; DD = developmental dyslexia; TD = typically developing.

#### Relationship between ERP and behavioural test performances

We further investigated whether the ERP responses correlated with reading performance in the DD group. The partial Pearson correlation analysis with age and IQ as covariates revealed that the P1 amplitude was negatively correlated with CCPM (*r* = -0.442, *p* = .039; **Figure 5A**). In other words, children with DD who had better passage reading fluency tended to show smaller P1 amplitude values during visual search. In addition, a significant positive correlation between the N2pc amplitude and reading comprehension was found in the DD group (*r* = 0.475, *p* = .026; **Figure 5B**). That is, children with DD who had better reading comprehension tended to show smaller N2pc amplitudes during visual search. The TD group showed a significant correlation between the N2pc amplitudes and behavioural performances in the visual search, including RT (*r* = 0.550, *p* = .001) and RTSD (*r* = 0.387, *p* = .026) (**Figure S1**). These results reveal that larger N2pc amplitudes predict better task performances in the TD group.

**Figure 5.**
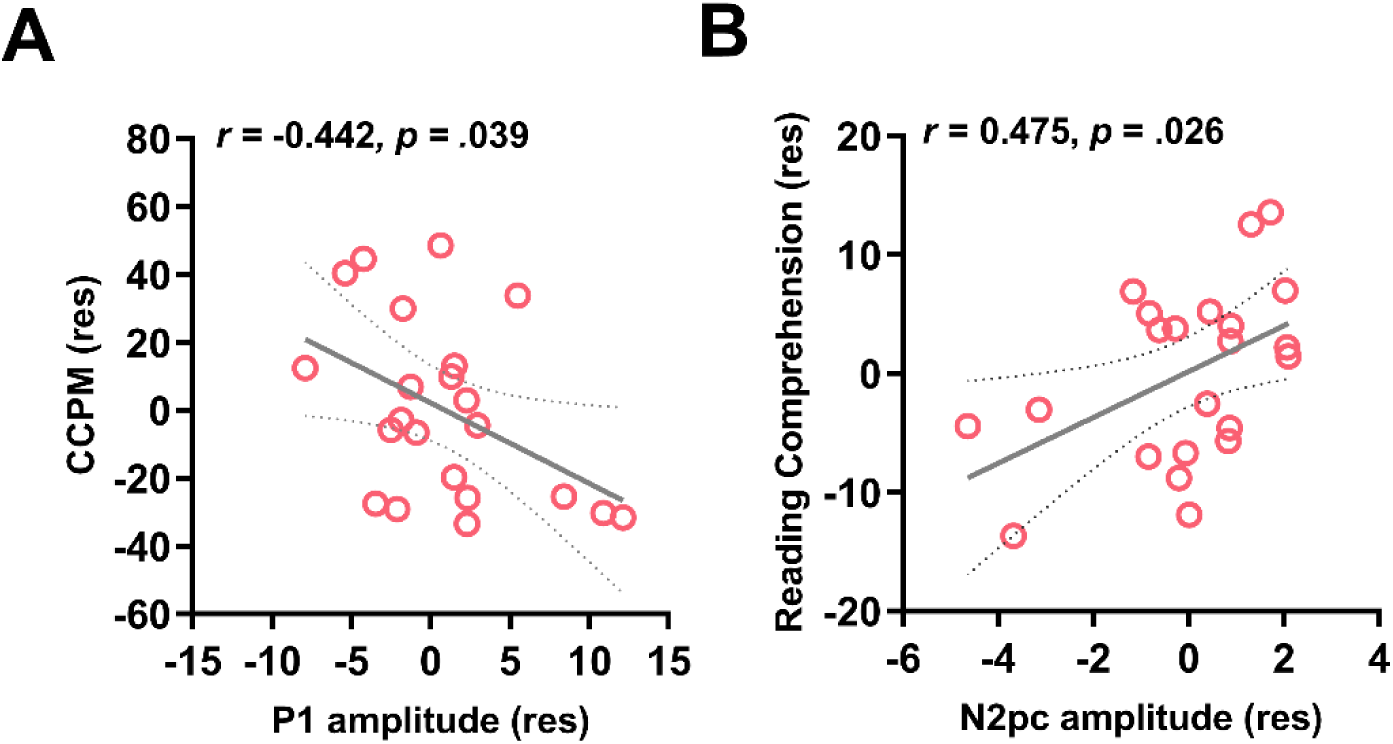
Partial correlation results between ERP components and behavioural performance in DD group. **(A)** Partial correlation between P1 amplitudes and CCPM. **(B)** Partial correlation between N2pc amplitudes and reading comprehension scores. Age and IQ were used as covariates to calculate the residual of each variable, and the residuals were used in each graph. CCPM = character read correctly per minute; DD = developmental dyslexia.

## Discussion

In this study, we investigated whether children with dyslexia exhibit impairments in visual selective attention. We applied a well-established pop-out search paradigm (Li et al., 2023; Sun et al., 2018; Wang et al., 2016) to capture children’s covert visual selective attention while recording EEG responses. N2pc and P_D_ components were used to assess selective attention and attentional suppression respectively. Although children with DD matched TD children in behavioural responses for both speed and accuracy, the EEG data revealed remarkable differences between the two groups. Compared with TD children, children with DD showed a larger and prolonged P1 component and the larger P1 amplitude was associated with lower reading fluency. Moreover, children with DD showed a larger target-evoked N2pc component which was associated with poor reading comprehension performance. More interestingly, target-elicited N2pc was followed by a positivity component (P_D_) in TD children but not in children with DD. The correlation between the N2pc and P_D_ implies that the absence of the P_D_ component in children with dyslexia may be related to their larger N2pc amplitudes.

The pop-out visual search task in our study tested visual selective attention to the target in DD and TD children. Interestingly, there were no significant differences in accuracy, RT, or RTSD between the DD and TD groups. This absence of a behavioural difference is consistent with previous findings (Iles et al., 2000; Koffman et al., 2023; Nguyen et al., 2021; Roach et al., 2007), suggesting that children with dyslexia can perform as well as TD children at the behavioural level. However, this does not imply that children with DD have intact visual selective attention as TD children do. Rather, our neural evidence reveals that they showed a different pattern of visual selective attention.

The electrophysiological results at the early sensory stage showed that both groups exhibited significant P1 components, reflecting early sensory processing and allocation of attentional resources to the stimuli at the sensory stage (Hillyard & Anllo, 1998; Luck, Heinze, Mangun, & Hillyard, 1990; for a review, see Koivisto & Revonsuo, 2010). Compared with TD children, we observed a larger amplitude and delayed latency of P1 in children with DD. Our results are consistent with previous findings in children with dyslexia, showing increased P1 amplitudes during words, visual and auditory processing (Csépe, Szücs, & Honbolygó, 2003; Kang, Lee, Park, & Leem, 2016; Silva, Ueki, Oliveira, Boggio, & Macedo, 2016) and slower neural responses (Kang et al., 2016; Meng, Liu, & Bi, 2022; Stefanics et al., 2011). Children with dyslexia exhibit a higher level of cortical excitability to recruit additional resources during early visual sensory processing, which may reflect the delayed development of synaptic pruning (Itier & Taylor, 2004; Taylor, Edmonds, McCarthy, & Allison, 2001). Prolonged P1 latency suggests a slower neural response and inefficient transmission of visual information (Taylor et al., 2001). Thus, increased effort (indexed by enhanced P1 amplitude) accompanied by a slower neural response (indexed by longer P1 latency) reflects the immaturity of visual processing in children with dyslexia at the early sensory stage.

The N2pc component is a widely accepted neurophysiological marker of selective attention. A previous study found that the N2pc amplitude was reduced in children compared to adults, reflecting insufficient attentional selection resources in children (Sun et al., 2018). In this study, both groups elicited a robust N2pc from the posterior region by the lateral target, as expected, reflecting successful selective attention to the target. For the first time, we found that children with dyslexia had significantly larger N2pc amplitudes, suggesting that they deployed more attentional resources towards the target. This result was also observed in a recent study by Koffman et al. (2023), in which the DD group showed a greater negative trend in the N2pc amplitude (Table 2 in Koffman et al., 2023). However, it was not statistically significant. This increase in N2pc may arise from insufficient engagement in early visual attention processing, for example, as indexed by P1, prompting them to compensate by allocating more selective attentional resources to the targets. Thus, the atypical selective attention in the DD group may indicate an attentional compensatory mechanism for dysfunctional early attentional processing.

The novel and critical point of our findings is the absence of the P_D_ component following N2pc in children with dyslexia, but a significant P_D_ in TD children. This lateral positivity elicited by the lateral target, also referred to as the P_T_ (Jannati et al., 2013; Sawaki et al., 2012), reflects the termination of target processing via attentional suppression. Children in the DD group elicited a larger N2pc but failed to trigger a subsequent P_D_ component, indicating that they not only needed to allocate more resources towards the target, but also struggled to disengage from the attended location and reorient to a new location. One possible explanation for the absence of the P_D_ component in the DD group could be their impaired ability in active attentional suppression, resulting in a failure to terminate attention from the current target and redirect it towards the subsequent target. However, this explanation requires further investigation, as the P_D_ component observed here was elicited by the target rather than the distractors. The robust positive correlation between N2pc and P_D_ only observed in the DD group provides a more plausible explanation for the absence of the P_D_ component, which is related to the atypical regulation of limited attentional resources. When excessive resources are devoted to attentional selection, fewer resources remain that children with DD could utilise for attentional disengagement, disturbing the balance in the regulation between attentional selection and inhibition. This N2pc/P_D_ relationship may provide insight into the neurological underpinnings of Sluggish Attentional Shifting (SAS) deficits in dyslexia (Hari, R., & Renvall, H., 2001). When dealing with rapidly changing stimulus sequences, individuals with dyslexia fail to smoothly disengage their attention from one item to another because of their inefficient attentional systems (Lallier et al., 2010). Thus, the overwhelming visual selective attentional resources and diminished attentional disengagement found in our study may contribute to a better understanding of sluggish attentional shifting in children with dyslexia.

We further demonstrated that increased P1 amplitude was negatively associated with CCPM. This aligns with previous research showing that better readers exhibit smaller P1 amplitudes (Wang et al., 2017). Therefore, such large P1 responses in children with DD point to an underlying problem in early visual processing that affects both reading performance and development (Maurer et al., 2007). More importantly, the smaller N2pc was associated with better reading comprehension performance in the DD group. Children with dyslexia who showed decreased N2pc amplitudes were more similar to TD children, potentially optimising resource allocation for target selection and disengagement. Previous studies using attentional training without phonological involvement improved reading ability in children with dyslexia (Franceschini et al., 2013; Wang et al., 2014). Therefore, attentional mechanisms play a vital role in visual word recognition and extraction by creating robust representations (Franceschini et al., 2022; Vidyasagar et al., 2010), that are essential steps in word comprehension (Lobier, Peyrin, Le Bas, & Valdois, 2012).

Overall, these findings draw a complex picture of visual attention ability in children with dyslexia, as they demonstrate immature early visual processing and excessive selective attentional resources to the target, but minimal attentional inhibition. As the reduced N2pc component reflects immature development (Sun et al., 2018) or impairments (Wang et al., 2016) of visual selective attention in children, hyperactivation of the N2pc and hypoactivation of P_D_ may indicate an atypical regulation of visual selective attention rather than a general deficit of visual attention in children with dyslexia. This distinct activation pattern may help explain why children with dyslexia struggle with tasks involving rapid or complex visual processing (Nguyen et al., 2021; Roach, & Hogben, 2007), particularly when the tasks have high cognitive demands, which make it harder for them to regulate their attention effectively. Therefore, we hypothesize that the atypical pattern of visual selective attention may be a crucial factor in the various visual attention deficits associated with dyslexia, although further studies are needed to confirm this hypothesis.

We excluded children with ADHD to eliminate the confusion regarding selective attention deficits caused by ADHD, as dyslexia and ADHD have been highly co-diagnosed (Willcutt & Pennington, 2000) and share overlapping symptoms such as learning and attentional challenges. Children with dyslexia were characterised by a larger N2pc followed by a diminished P_D_ component, while children with ADHD were characterised by a smaller N2pc and a diminished P_D_ component (Wang et al., 2016). Our results are consistent with the notion that DD and ADHD are distinct disorders that exhibit different patterns of visual selective attention in behavioural (for a review, see Hokken et al., 2023) and neural responses.

We admit that some TD children did not complete all the reading tests. Based on the questionnaire, TD children achieved excellent scores on school reading tests and reported no difficulties in learning Chinese. In future studies, tasks with more salient distractors will enable the investigation of both visual selective attention and active suppression in children with dyslexia, especially in the distractor-elicited P_D_ component.

## Conclusion

This study provides novel and strong neurophysiological evidence that children with dyslexia have deficits in visual selective attention, which might be related to their atypical regulation of attentional resources and associated with their reading abilities. The correlation between N2pc and P_D_ suggests an imbalance between attentional selection, indexed by the larger N2pc and attentional suppression, indexed by the absence of P_D_, in children with dyslexia. These findings shed new light on the atypical visual selective attention in dyslexia and offer a new perspective for exploring the underlying mechanism of visual attention deficits in children with dyslexia.

## Acknowledgements

This work was supported by the STI 2030–Major Projects (2021ZD0200500) and the National Natural Sciences Foundation of China (NSFC: 32371141). We are especially grateful to the children and students who participated in this study.

## Ethical standards

The authors assert that all procedures contributing to this work comply with the ethical standards set by the relevant national and institutional committees on human experimentation and with the Helsinki Declaration of 1975, as revised in 2008.

## Data Availability Statement

The data that support the findings of this study are available on request from the corresponding author.

