## Supplemental Figure for "Atypical Visual Selective Attention in Children with Dyslexia: Evidence from N2pc and PD"

### *Supplement*

#### Supplemental Figure

#### Figure S1

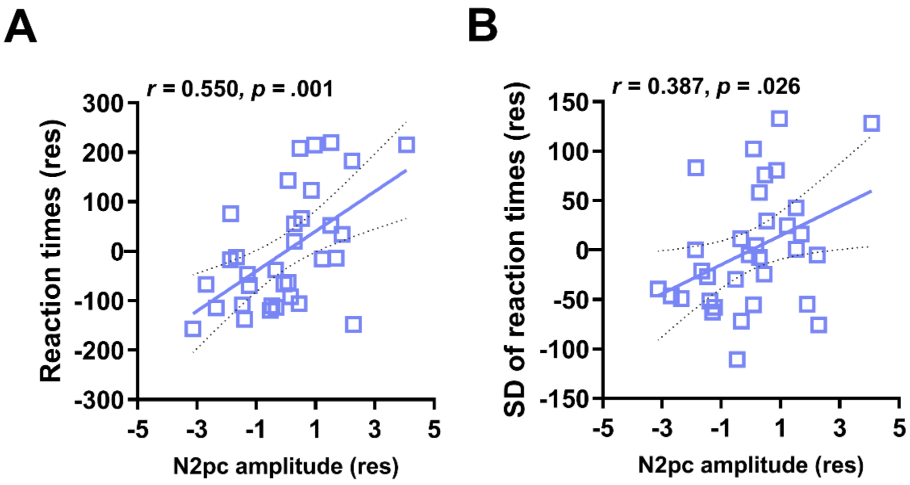


**Figure S1.** Partial correlation results between N2pc amplitudes and behavioural performance in TD group. **(A)** Partial correlation between N2pc amplitudes and reaction times. **(B)** Partial correlation between N2pc amplitudes and SD of reaction times. Age and IQ were used as covariates to calculate the residual of each variable, and the residuals were used in each graph. SD= standard deviation; TD = typically developing.
